# Establishment of a second-generation transgenic marmoset model of polyglutamine disease recapitulating neurological symptoms and pathology

**DOI:** 10.1101/2025.03.12.642922

**Authors:** Eiko N. Minakawa, Kensuke Owari, Naotake Nogami, Terumi Nakatani, Masashi Koizumi, Akiyo Kawanobe, Yosuke Saga, Moeko Kudo, Satoru Noguchi, Takashi Hanakawa, Yuki Hori, Hidemi Numazawa, Toshihide Takeuchi, Yuko Katakai, Ryoichi Saito, Yoshitaka Nagai, Yuko Saito, Ikuo Tomioka, Kazuhiko Seki

**Affiliations:** Department of Neurophysiology, National Institute of Neuroscience, National Center of Neurology and Psychiatry, Kodaira, Tokyo, Japan; The Corporation for Production and Research of Laboratory Primates, Tsukuba, Ibaragi, Japan; Department of Neuromuscular Research, National Institute of Neuroscience, National Center of Neurology and Psychiatry, Kodaira, Tokyo, Japan; Integrative Brain Imaging Center, National Center of Neurology and Psychiatry, Kodaira, Tokyo, Japan; Department of Integrated Neuroanatomy and Neuroimaging, Kyoto University Graduate School of Medicine, Sakyo, Kyoto, Japan; Life Science Research Institute, Kindai University, Osaka-Sayama, Osaka, Japan; Administrative Section of Primate Research Facility, National Institute of Neuroscience, National Center of Neurology and Psychiatry, Kodaira, Tokyo, Japan; Department of Neurology, Kindai University Faculty of Medicine, Osaka-Sayama, Osaka, Japan; Department of Pathology and Laboratory Medicine, National Center of Neurology and Psychiatry Hospital, Kodaira, Tokyo, Japan; Laboratory of Applied Reproductive Science, Faculty of Agriculture, Shinshu University, Kami-Ina, Nagano, Japan

**Keywords:** Common marmoset, polyglutamine diseases, spinocerebellar ataxia 3, Machado-Joseph disease, germline transmission, home-cage behavior assessment

## Abstract

Neurodegenerative diseases, including polyglutamine diseases, remain a major clinical challenge, partly because of limited animal models that recapitulate human disease. Here, we describe a second-generation transgenic marmoset model of spinocerebellar ataxia 3 (SCA3), a polyglutamine disease, which stably expresses expanded CAG repeats in *ATXN3*. All five offspring of the founder marmoset harbored the transgene with reduced transgene integration sites and without repeat instability or genetic mosaicism, offering improved construct validity. Three of the five marmosets developed progressive motor impairments that segregated into two distinct phenotypes: early onset with rapid progression and late onset with mild progression, accompanied by corresponding patterns in body weight gain and grip strength. Pathological analysis revealed cerebellar Purkinje cell loss, spinal cord neurodegeneration, and widespread intranuclear inclusions. The severity of motor phenotypes correlated with transgene expression levels in disease-relevant brain regions, including the cerebellum, spinal cord, and striatum. By overcoming the common translational limitations of rodent systems, our second-generation model offers a powerful platform for investigating disease mechanisms and testing potential therapeutic interventions. Our results advance the utility of transgenic marmosets as clinically relevant models of neurodegenerative diseases.

**Summary Statement:** Second-generation transgenic marmoset models of spinocerebellar ataxia 3 replicated the progressive motor deficits and neuropathology of the founder marmoset, providing a powerful platform for studying disease mechanisms and developing therapies.

## Introduction

Neurodegenerative diseases, such as Alzheimer’s disease, Parkinson’s disease, amyotrophic lateral sclerosis, and polyglutamine (polyQ) diseases, are a group of disorders characterized by the progressive and selective loss of neuronal cells in various regions within the central nervous system (CNS) due to abnormal neurotoxic protein accumulation that are specific to each disease (Nagai and Minakawa, 2015; Wilson et al., 2023). Despite substantial advancements in our understanding of the underlying pathomechanisms of these diseases, there is a near absence of efficacious and accessible disease-modifying therapies that can slow or halt their progression, representing an important unmet clinical need (Belder et al., 2023; Kalia et al., 2024; Kiernan et al., 2021; Malik et al., 2021).

Furthermore, most of these diseases lack specific biomarkers, particularly during the early phases when neuronal loss is minimal and disease-modifying interventions would be most effective (Hansson, 2021).

Mouse models of neurodegenerative diseases have been used extensively to develop diagnostic biomarkers and therapeutic agents for these diseases. However, genetic, anatomical, and physiological differences between rodents and humans have made it difficult to translate preclinical findings from rodents to clinical applications in humans (McGonigle and Ruggeri, 2014). In contrast, non-human primates (NHPs), such as rhesus macaque (*Macaca mulatta*), cynomolgus macaque (*M. fascicularis*), and common marmoset (*Callithrix jacchus*), share highly similar genetic, anatomical, physiological, and behavioral traits with humans. They also have a larger brain size and longer lifespan than rodents, making them valuable experimental animals for modeling human diseases (Chan, 2013). NHP models of neuronal diseases are thus expected to enhance our understanding of neurological and psychiatric disorders, including neurodegenerative diseases (Belmonte et al., 2015).

Recent technological advance for introducing genetic modification into NHPs such as common marmosets and rhesus or cynomolgus macaques have allowed for the development of improved NHP models of human diseases including neurodegenerative diseases (Chan, 2013; Liu et al., 2016; Schmidt et al., 2022). Among these NHPs, common marmosets offer many advantages for rapidly generating gene-modified disease models, such as smaller size, faster sexual maturation, shorter gestation period, and higher fecundity than rhesus of cynomolgus macaques (Tomioka et al., 2018). Technologies for generating transgenic marmosets with germline transmission were first established by Sasaki and colleagues (Sasaki et al., 2009). We previously utilized these techniques and established a transgenic marmoset model of the polyQ disease spinocerebellar ataxia 3 (SCA3) / Machado-Joseph disease, which is the most common subtype of autosomal dominant spinocerebellar ataxia (Tomioka et al., 2017). Our transgenic SCA3 marmoset model recapitulated the human disease process including the neuronal accumulation of abnormal polyQ proteins and neuronal cell loss, resulting in neurological symptoms such as motor impairment (Tomioka et al., 2017). We also confirmed the germline transmission of the transgene and successfully produced second-generation offspring of the transgenic marmosets (Tomioka et al., 2017). In the present study, we produced further second-generation offspring marmosets and assessed whether the genetic, behavioral, and pathological phenotypes of the founder marmoset were transmitted to these second-generation marmosets towards establishing a stable polyQ disease model line for future colony expansion.

## Results

### Production of second-generation transgenic marmosets expressing expanded CAG repeats

We produced second-generation marmosets from one symptomatic founder marmoset (PQD1), one of five founder transgenic marmosets carrying CMV-*ATXN3*-120Q-IRES-Venus generated in our previous study (Tomioka et al., 2017). We chose PQD1 because this was the only male animal that showed moderate disease progression and reached sexual maturity (Tomioka et al., 2017). We injected the ejaculated sperm of PQD1 into the oocytes of wild-type marmoset by intracytoplasmic sperm injection and transferred 69 embryos into 29 surrogate mothers. Six surrogates became pregnant and five surrogates delivered seven offspring. Among them, two marmosets were stillborn.

Five marmosets (PQD11-15) were thus obtained; PQD13 and 14 were littermates. In our previous work, we analyzed and described PQD11–14; PQD15 was born after this earlier report.

### Genetic analyses of the second-generation marmosets

We first analyzed whether the transgene of the founder marmoset (Fig. 1A) was transmitted to the second-generation marmosets. Polymerase chain reaction (PCR) analysis of genomic DNA or complementary DNA (cDNA) obtained from skin-derived fibroblast cell lines established from each second-generation marmoset demonstrated that all five marmosets carried and expressed the *ATXN3*-120Q transgene (Fig. 1B). We then compared the genetic characteristics of the second-generation marmosets and the founder marmoset. CAG repeat analysis using the skin-derived fibroblast cell lines showed that all five second-generation marmosets expressed the *ATXN3*-120Q transgene with a CAG repeat length that was identical to that of the founder (Fig. 1C). Quantitative real-time PCR analyses using the skin-derived fibroblasts revealed that the transgene expression levels of all second-generation marmosets except PQD12 were comparable to that of PQD1 (Fig. 1D). Lentiviral integration site analysis by genome walking demonstrated that two, three, or four of the six integration sites in PQD1 was transmitted to PQD11–15 (Fig. 1E). Copy number analysis of the integrated transgene using digital PCR revealed that two, three, or four copies of transgenes were integrated into PQD11–15 (Fig. 1F). The comparison of the number of integration sites (Fig. 1E) and the copy number of the transgene (Fig. 1F) within each marmoset suggested that one copy of the transgene was integrated into each integration site in PQD11, 12, and 15, whereas two copies were integrated into either of the integration sites in PQD13 and 14. Alternatively, the integration site of one of the three copies of the transgene may not have been detected by genome walking in PQD13 and 14. Overall, these results indicate that the genetic characteristics of the founder marmoset were successfully inherited by the second-generation marmosets.

**Figure 1.**
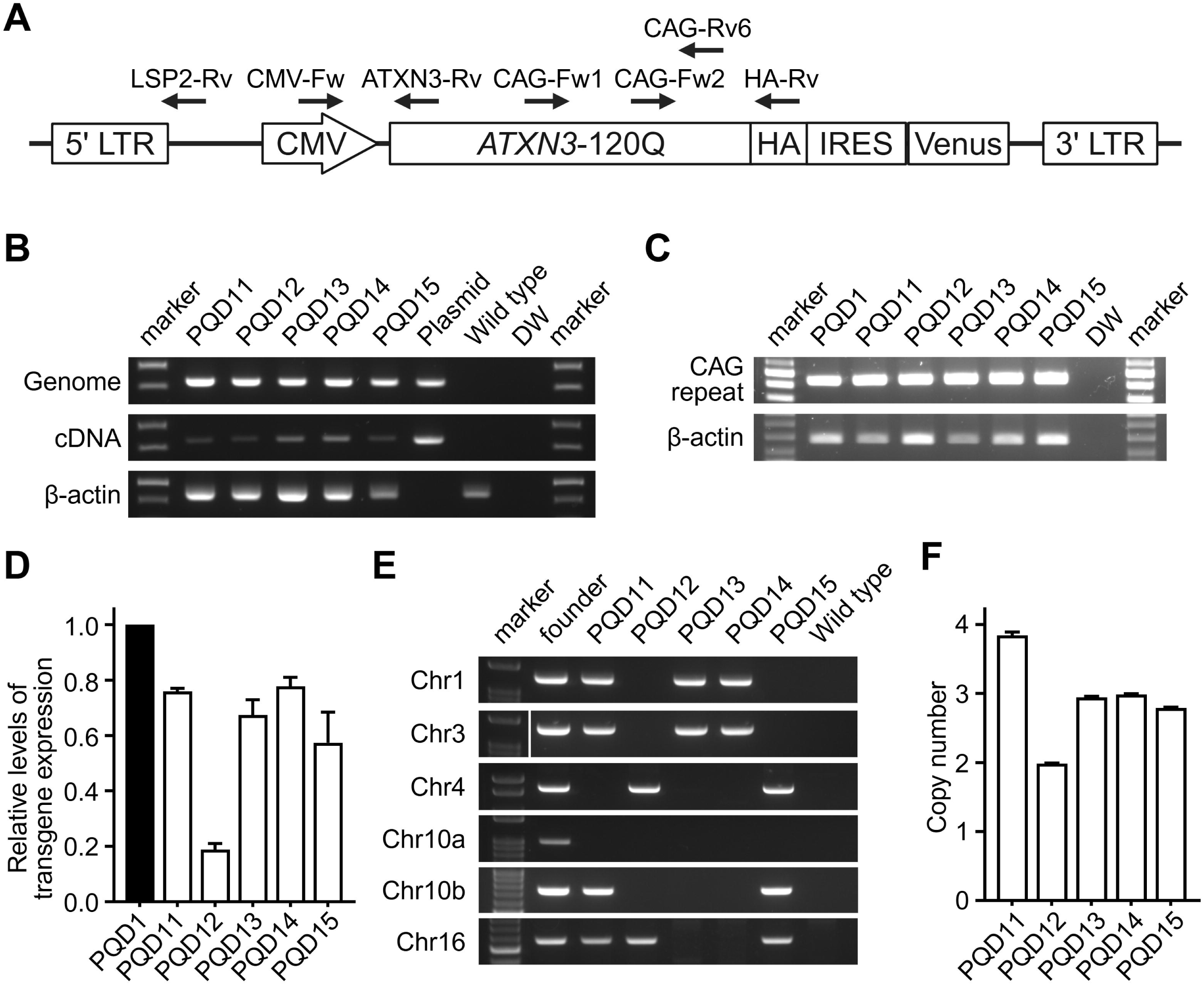
Generation of second-generation ataxin 3-120Q transgenic marmosets. **A**. Schematic representation of the genetic construct introduced into the marmoset embryos to generate the founder marmoset PQD1 (Tomioka et al., 2017). Black arrows indicate the primers used for genomic polymerase chain reaction (PCR), reverse-transcription PCR (RT-PCR), quantitative RT-PCR, integration site analysis, and copy number analysis in this study. **B**. Representative images of the genomic PCR (top) and RT-PCR (middle) analyses of the second-generation marmosets (PQD11–15), a positive control (plasmid), and negative controls (wild type and water; DW). β-actin (bottom) was used as an internal control. **C**. Representative image of the CAG repeat analysis (top) of the founder marmoset (PQD1), the second-generation marmosets (PQD11–15), and a negative control (water; DW). β-actin (bottom) was used as an internal control. **D**. Relative transgene expression levels of the founder marmoset (PQD1) and the second-generation marmosets (PQD11– 15). Transgene expression was normalized using GAPDH, and the normalized transgene expression level of PQD1 was set as 1. **E**. Representative image of the transgene integration site analysis of the founder marmoset (PQD1), the second-generation marmosets (PQD11–15), and negative controls (wild type and water). **F**. Bar graph showing the transgene copy number of the second-generation marmosets (PQD11–15). In **D** and **F**, data are presented as the mean ± s.e.m. of four and two independent experiments, respectively. Each dot represents data from a single experiment.

### *In vivo* phenotype analyses of the second-generation marmosets; behavioral assessments

We then analyzed the *in vivo* phenotypes of the second-generation marmosets focusing on their natural behaviors in the home cage. The quantitative detection of home-cage behavior enables prolonged and unbiased observations of spontaneous behavior in a familiar environment for the animals, thus providing a reproducible measure of animal behaviors (Grieco et al., 2021). We developed a semiquantitative rating scale (the Marmoset Behavioral Assessment Scale [MBAS]; Table 1) to longitudinally assess the motor function of the second-generation marmosets using video of their home-cage behavior. We assessed five different aspects of movements, namely, posture, activity, motility, muscle strength, and balance, in their natural behavior; these aspects were scored by a rater who was blinded to the timing of each video recording.

**Table 1.** Marmoset Behavioral Assessment Scale (MBAS).

| Score | Posture | Activity (Frequency of spontaneous movements) | Motility (Smoothness of spontaneous movements) | Muscle strength | Balance (Frequency of slipping and falling) |
| --- | --- | --- | --- | --- | --- |
| 0 | Normal | Normal | Normal | Normal | Normal |
| 1 | Mildly abnormal (e.g. hunched posture) | Mildly decreased | Mildly decreased (e.g. mildly clumsy movements) | Mildly decreased (e.g. occasional slipping when hanging on cage wires) | Mildly imbalanced (e.g. slipping on the perch or the loft) |
| 2 | Moderately abnormal (e.g. mostly leans against the wall, difficult to stand) | Moderately decreased (e.g. mostly stays in one place) | Moderately decreased (e.g. irregular and incoordinated movements) | Moderately decreased (e.g. frequent slipping when hanging on cage wires) | Moderately imbalanced (e.g. frequent slipping on the cage floor) |
| 3 | Unmeasurable (e.g. mostly lies on the cage floor) | Severely decreased | Severely decreased | Severely decreased | Severely imbalanced or unmeasurable (e.g. mostly lies on the cage floor) |

MBAS scoring demonstrated that PQD13 and 14 exhibited an earlier onset of motor impairment, at around 19 weeks of age, than that of PQD11, at around 27 weeks of age (Fig. 2A). In addition, the early-onset group (PQD13 and 14) showed a faster progression of motor impairment than the late onset group (PQD11) (Fig. 2A). To confirm these behavioral changes, we measured the in-cage motility of the second-generation marmosets using an infrared motion sensor placed above the cage (Koizumi et al., 2021). The total activity count per day demonstrated an acute decline following the early onset of motor impairment in PQD13 and 14 and a gradual decline following the later onset of motor impairment in PQD11 (Fig. 2B). These changes were in line with the changes in MBAS scores (Fig. 2A). MBAS scores indeed showed a statistically significant positive correlation with activity counts (r =-0.9467, p < 0.0001; Fig. 2C). Collectively, these results indicate that the MBAS is a useful method for detecting motor impairments in marmosets because it has a comparable sensitivity with continuous quantitative activity measurement using an infrared sensor but does not require special devices. Importantly, these results also indicate that our second-generation marmosets can be categorized into three different groups: symptomatic with early onset and rapid progression of motor impairments (PQD13 and 14), symptomatic with later onset and slower progression of motor impairments (PQD11), and asymptomatic (PQD12 and 15).

**Figure 2.**
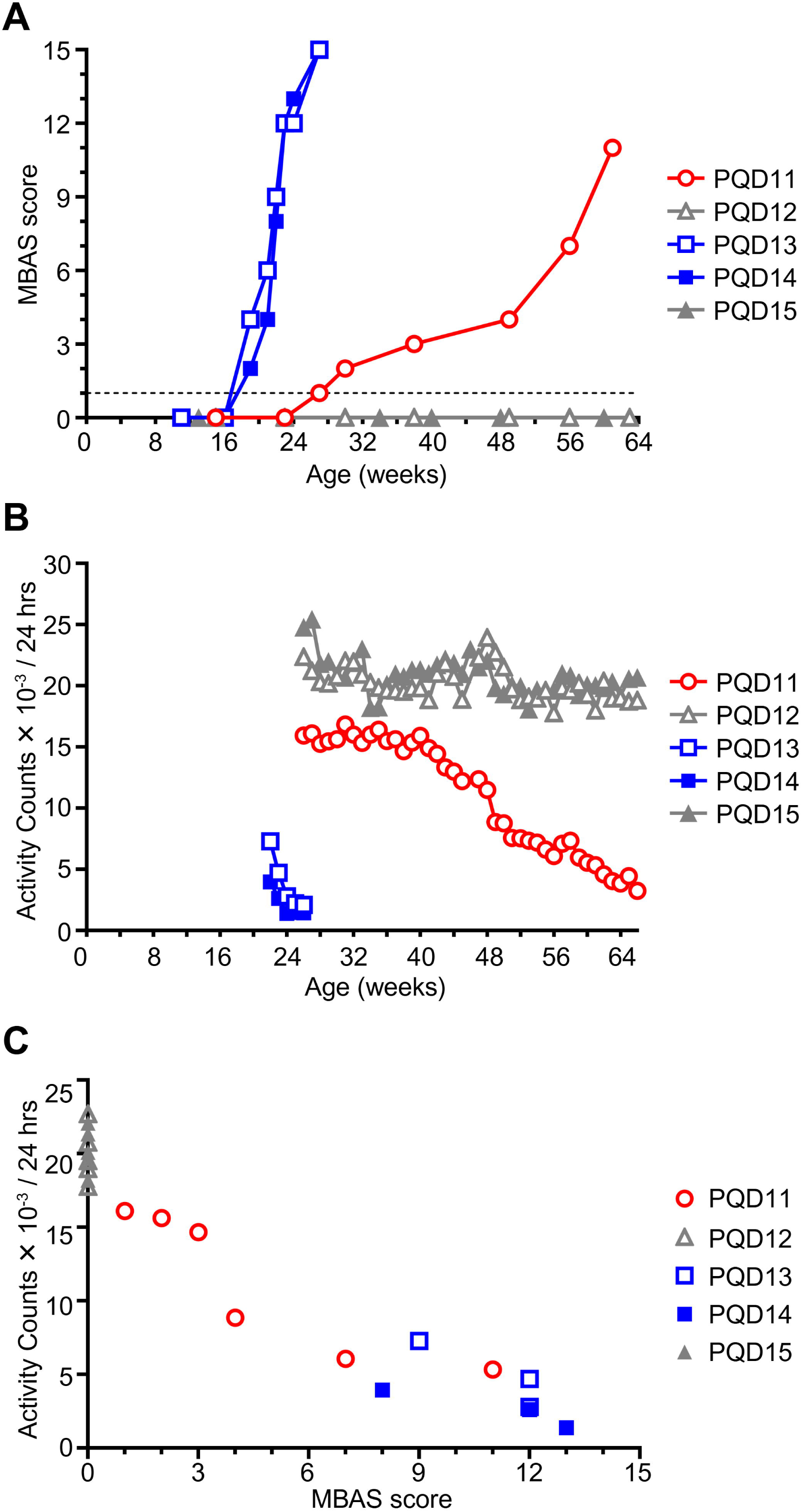
Second-generation transgenic marmosets show progressive motor impairment. **A.** Longitudinal assessment of the motor function of the second-generation marmosets (PQD11–15) using the Marmoset Behavioral Assessment Scale (MBAS). **B.** Longitudinal assessment of the motor function of the second-generation marmosets (PQD11–15) using infrared motion sensors placed above the home cages. **C.** Scatter plot showing the correlation between the motor function assessments using the MBAS and infrared motion sensors (r = - 0.9467, p < 0.0001; Spearman’s correlation analysis). In **A**–**C**, data from symptomatic marmosets with early onset and rapid progression (PQD13 and 14), symptomatic marmoset with later onset and slower progression (PQD11), and asymptomatic marmosets (PQD12 and 15) are presented in blue, red, and gray, respectively.

### Comparison of *in vivo* phenotypes between the founder and second-generation marmosets

Next, we compared the *in vivo* phenotypes of the first-generation (founder) and second-generation marmosets by comparing the phenotypes that were available for both groups: body weight and grip strength. Longitudinal body weight measurement showed that the founder marmoset, PQD1, gained almost no body weight after the onset of motor impairments at 3–4 months of age, as described previously (Figs. 3A, B; Tomioka et al., 2017). Among the second-generation marmosets, the two early-onset marmosets, PQD13 and 14, showed an acute decline in body weight gain starting at 14–15 weeks of age, which was earlier than that of PQD1; they gained almost no body weight thereafter, similar to PQD1 (Figs. 3A and B). In contrast, the late-onset marmoset, PQD11 showed a gradual decline in body weight gain starting at around 24 weeks of age and gained almost no body weight after around 40 weeks of age (Fig. 3A). The body weight gain of the two asymptomatic marmosets, PQD 12 and 15, remained equivalent to that of the wild-type marmosets throughout the observation period (Fig 3A). These results support our observations from the behavioral assessment that the symptomatic marmosets can be categorized into two groups: early-and later onset. Next, we compared the body weight gain per week before and after symptom onset separately for the marmosets with early-(Fig. 3C) and late (Fig. 3D) onset. In accord with the body weight changes in PQD1 before and after motor impairment onsets (Tomioka et al., 2017), after behavioral abnormality onset was detected using the MBAS, the weekly body weight gain of all symptomatic second-generation marmosets (PQD13 and 14 in Fig. 3C, and PQD11 in Fig. 3D) was significantly lower than that of age-matched asymptomatic marmosets (PQD12 and 15) and wild-type marmosets.

**Figure 3.**
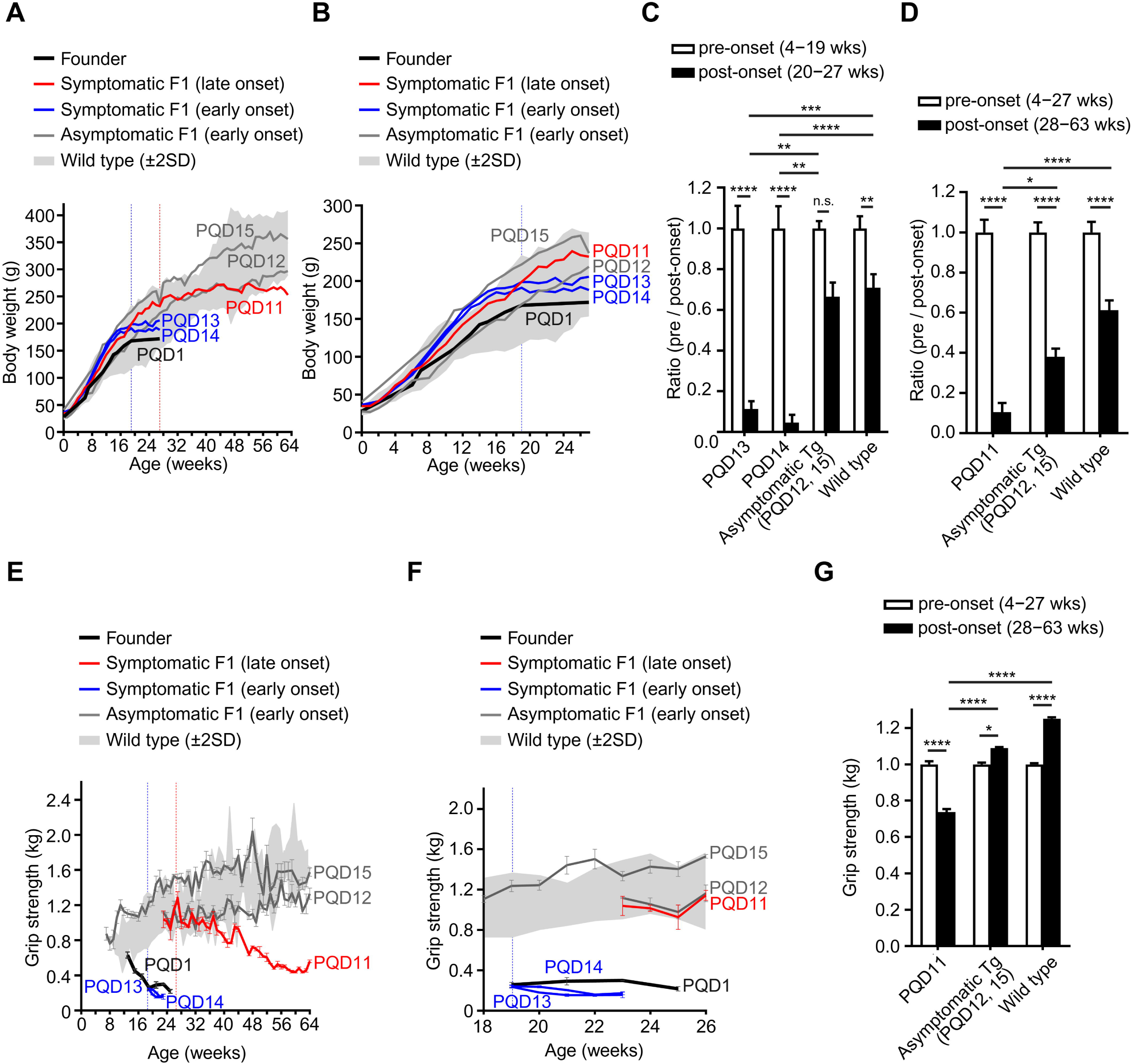
Comparisons of the physical and behavioral phenotypes between the founder and second-generation marmosets. A and. **B.** Longitudinal assessment of the body weights of the founder marmoset (PQD1), second-generation marmosets (PQD11–15), and wild-type marmosets. Expanded view of **A** during the first 27 weeks of age is shown in **B**. Data from symptomatic marmosets with early onset and rapid progression (PQD13 and 14), the symptomatic marmoset with later onset and slower progression (PQD11), and asymptomatic marmosets (PQD12 and 15) are presented as blue, red, and gray lines, respectively. The gray shaded area represents the mean ± s.e.m. of the wild-type marmosets (n = 11). Vertical red or blue dotted lines indicate the age of onset of motor impairment detected using the MBAS in marmosets with early onset and rapid progression (PQD13 and 14) or late onset and slow progression (PQD11), respectively. **C and D.** Bar graph showing the comparison of weekly body weight gain before and after the onset of motor impairment in second-generation symptomatic marmosets with early onset and rapid progression (**C**; PQD13 and 14) and the symptomatic marmoset with late onset and slower progression (**D**; PQD11). Age-matched data from asymptomatic marmosets (PQD12 and 15) and wild-type marmosets are shown as controls. Data are presented as the mean ± s.e.m. of the ratio of weekly body weight gain before and after the age of symptomatic onset in the early-onset marmosets (**C**; PQD13 and 14) and late-onset marmoset (**D**; PQD11). The pre-onset mean of weekly body weight gain in each group was set as 1. **E and F.** Longitudinal assessment of the grip strength of the founder marmoset (PQD1), second-generation marmosets (PQD11–15), and wild-type marmosets. Expanded view of **E** from 18 to 26 weeks of age is shown in **F**. Data from symptomatic marmosets with early onset and rapid progression (PQD13 and 14), the symptomatic marmoset with later onset and slower progression (PQD11), and asymptomatic marmosets (PQD12 and 15) are presented as blue, red, and gray lines, respectively. The gray shaded area represents the mean ± s.e.m. of wild-type marmosets (n = 5). Vertical red or blue dotted lines indicate the age of onset of motor impairment detected using the MBAS in marmosets with early onset and rapid progression (PQD13 and 14) or late onset and slow progression (PQD11), respectively. Data are shown as the mean ± s.e.m. of the five measurements at each timepoint. **G.** Bar graph showing the comparison of grip strengths before and after the onset of motor impairment in **the** second-generation symptomatic marmoset with late onset and slower progression (**D**; PQD11). Age-matched data of asymptomatic marmosets (PQD12 and 15) and wild-type marmosets are shown as controls. Data are presented as the mean ± s.e.m. of the ratio of grip strengths before and after the age of symptomatic onset in the late-onset marmoset (**D**; PQD11). The pre-onset mean of grip strength in each group was set as 1. In **C**, **D**, and **G**, statistical analyses were performed to assess the differences between the ratios of pre-and post-onset phenotypes using two-way ANOVA followed by Tukey’s multiple comparison test (* p < 0.05, ** p < 0.01, *** p < 0.001, **** p < 0.0001, n.s. not significant).

Intriguingly, considering that the onset of motor impairments detected using the MBAS was at around 19 (for PQD13 and 14) or 27 (for PQD11) weeks of age (Figs. 2A and B), the onset of the decline in body weight gain was a sensitive predictor of future motor impairment starting 3–4 weeks before motor impairment onset in both early-and late-onset symptomatic marmosets.

We next analyzed changes in grip strength. In accord with the findings of body weight changes, the early-onset marmosets (PQD13 and 14) showed significantly weaker grip strength than wild-type marmosets at 19–23 weeks of age; their grip strength was as weak as that of PQD1 (Figs. 3E, F). In contrast, the grip strength of the late-onset marmoset (PQD11) showed a gradual and milder decline compared with that of PQD1, 13 and 14 (Fig. 3E) but was significantly weaker than that of asymptomatic (PQD12 and 15) and wild-type marmosets (Fig. 3G).

Together, these results indicate that, among the symptomatic second-generation marmosets, marmosets with early onset exhibited physical and behavioral phenotypes equivalent to those of the founder marmoset, whereas the marmoset with later onset exhibited milder phenotypes.

### Pathological analyses of symptomatic second-generation marmosets

We then analyzed the pathological findings of the symptomatic marmosets and compared them with those of PQD1. We previously reported that the founder marmoset (PQD1) showed marked neurodegeneration in the cerebellum and spinal cord and exhibited intranuclear inclusions containing the expanded polyQ stretch in the cerebrum, brainstem, and spinal cord (Table 2; Tomioka et al., 2017). PQD1 also showed pathologies in the peripheral nerves and skeletal muscles, such as acute axonal degeneration and chronic degeneration of the myofibers; these were also accompanied by intranuclear inclusions (Table 2; Tomioka et al., 2017).

**Table 2.** Summary of the neuropathological findings of the founder (PQD1) and symptomatic second-seneration (PQD11, 13, and 14) marmosets.

| Degeneration | PQD1 | PQD11 | PQD13 | PQD14 |
| --- | --- | --- | --- | --- |
| Cerebrum | – | ± | – | – |
| Cerebellum | ++ | + | + | ± |
| Brain stem | – | – | – | – |
| Spinal cord | ++ | ++ | ++ | ++ |
| Peripheral nerves | ++ | ++ | N/E | N/E |
| Skeletal muscles | +++ | ++ | +++ | +++ |

| Inclusions | PQD1 | PQD11 | PQD13 | PQD14 |
| --- | --- | --- | --- | --- |
| Cerebrum | ± | + | ± | + |
| Cerebellum | – | – | – | – |
| Brain stem | ± | ± | ± | ± |
| Spinal cord | ++ | ++ | ++ | ++ |
| Peripheral nerves | N/E | + | N/E | N/E |
| Skeletal muscles | ++ | +++ | +++ | +++ |

N/E, not examined

Among the symptomatic second-generation marmosets, PQD11, the late-onset marmoset, exhibited relatively mild loss of Purkinje cells in the cerebellum (Fig. 4A), accompanied by mild gliosis (Fig. 4B) without intranuclear inclusions. In the spinal cord, PQD11 showed marked neurodegeneration (Figs. 4C–E) and abundant eosinophilic intranuclear inclusions (Fig. 4E) that were immunoreactive for both the expanded polyQ stretch (Fig. 4F) and ubiquitin (Figs. 4G, H).

**Figure 4.**
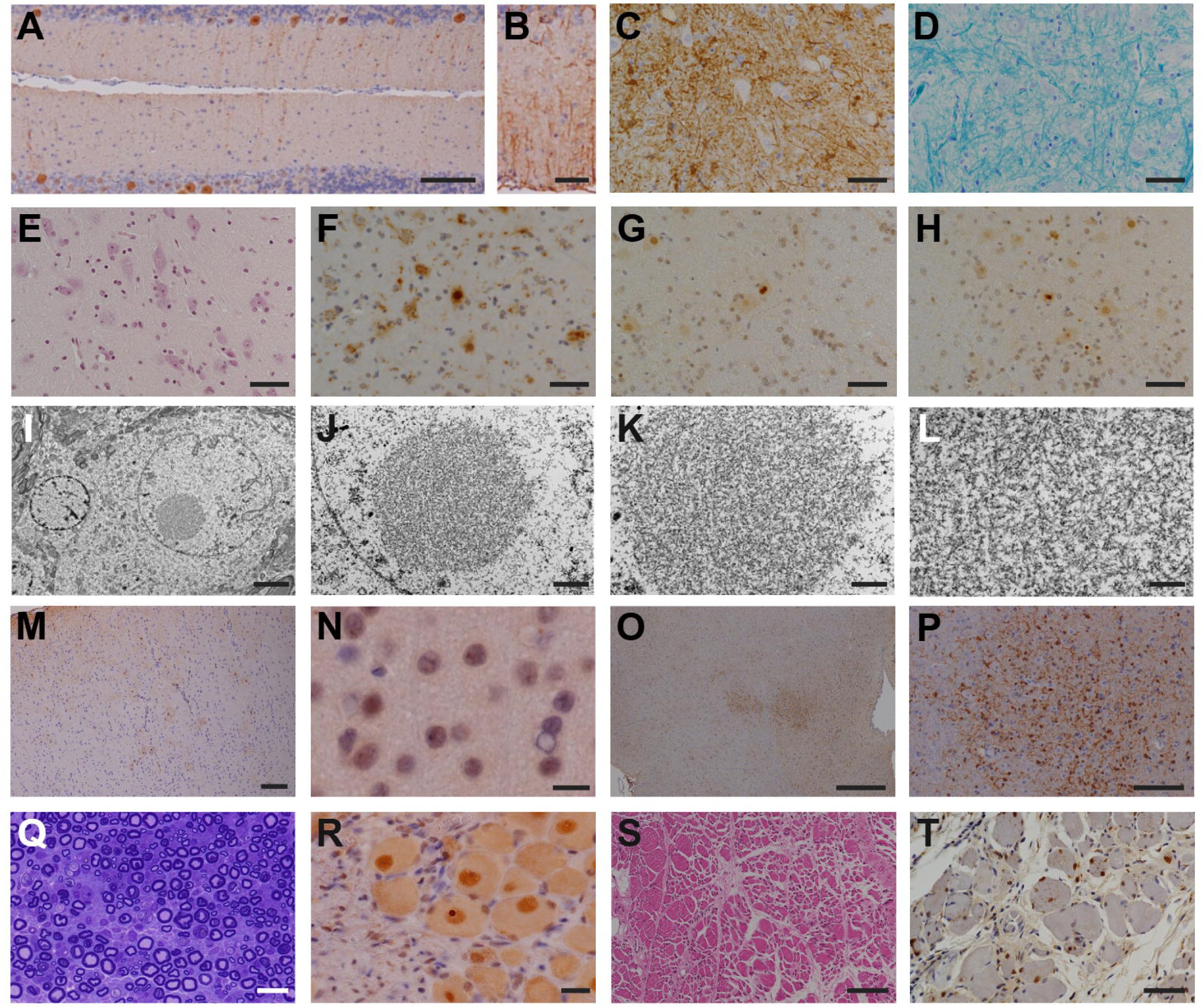
Degeneration and inclusion body formation in the brain, spinal cord, peripheral nerve and skeletal muscle of PQD11. A and. **B.** Anti-calbindin (**A**) and anti-GFAP (**B**) staining of the cerebellum of PQD11. **C–H.** Anti-GFAP (**C**), Klüver-Barrera (**D**), hematoxylin and eosin (**E**), anti-1C2 (**F**), and anti-ubiquitin (**G** and **H**) staining of the lumbar spinal cord of PQD11. **I–L**. Ultrastructural images of the inclusion body in the lumbar spinal cord of PQD11. **M and N.** Anti-GFAP (**M**) and anti-ubiquitin (**N**) staining of the cerebrum of PQD11. **O and P.** Anti-Iba1 staining of the brainstem of PQD11. A magnified image of the dotted square in **O** is shown in **P**. **Q.** Toluidine blue staining of the peripheral nerve of PQD11. **R.** Anti-ubiquitin staining of the peripheral nerve (dorsal root ganglion) of PQD11. **S and T.** Hematoxylin and eosin (**S**) and anti-1C2 (**T**) staining of the skeletal muscle (quadriceps femoris) of PQD11. Scale bars: 100 μm in A, M, P, and S; 50 μm in B–H and T; 5 μm in I; 2 μm in J; 1 μm in K; 500 μm in L and O; 10 μm in N; 20 μm in Q–R.

Nuclear inclusion bodies are circular structures of approximately 2-5 microns, with no limiting membrane, and are formed by randomly arranged electron-dense fibers of 12-15 nm diameter (Figs. 4I–L). Furthermore, PQD11 showed mild neurodegeneration in the cerebrum (Fig. 4M) accompanied by scarce intranuclear inclusions (Fig. 4N) and neurodegeneration without intranuclear inclusions was observed in the brainstem (Figs. 4O–P).

In addition, PQD11 exhibited acute axonal degeneration in the peripheral nerve of the lower limb (Fig. 4Q) with ubiquitin-positive inclusions (Fig. 4R). PQD1 also showed degenerating myofibers, suggesting chronic myopathic changes (Fig. 4S), accompanied by intranuclear inclusions that were immunoreactive for the expanded polyQ stretch (Fig. 4T) in the skeletal muscles of the lower limb.

By contrast, PQD13 and 14, the early-onset marmosets, showed a loss of Purkinje cells in the cerebellum without intranuclear inclusions (Fig. 5A, H) as well as marked neurodegeneration in the spinal cord (Fig. 5B, I) accompanied by abundant intranuclear inclusions (Fig. 5C, D, and J). In addition, both PQD13 and 14 showed intranuclear inclusions in the cerebrum (Fig. 5E, K). No neurodegeneration was observed in the brainstems of PQD13 or 14 (Fig. 5F, L); however, scarce intranuclear inclusions were observed in the oculomotor nuclei of PQD13 and 14 (Fig. 5G, M).

**Figure 5.**
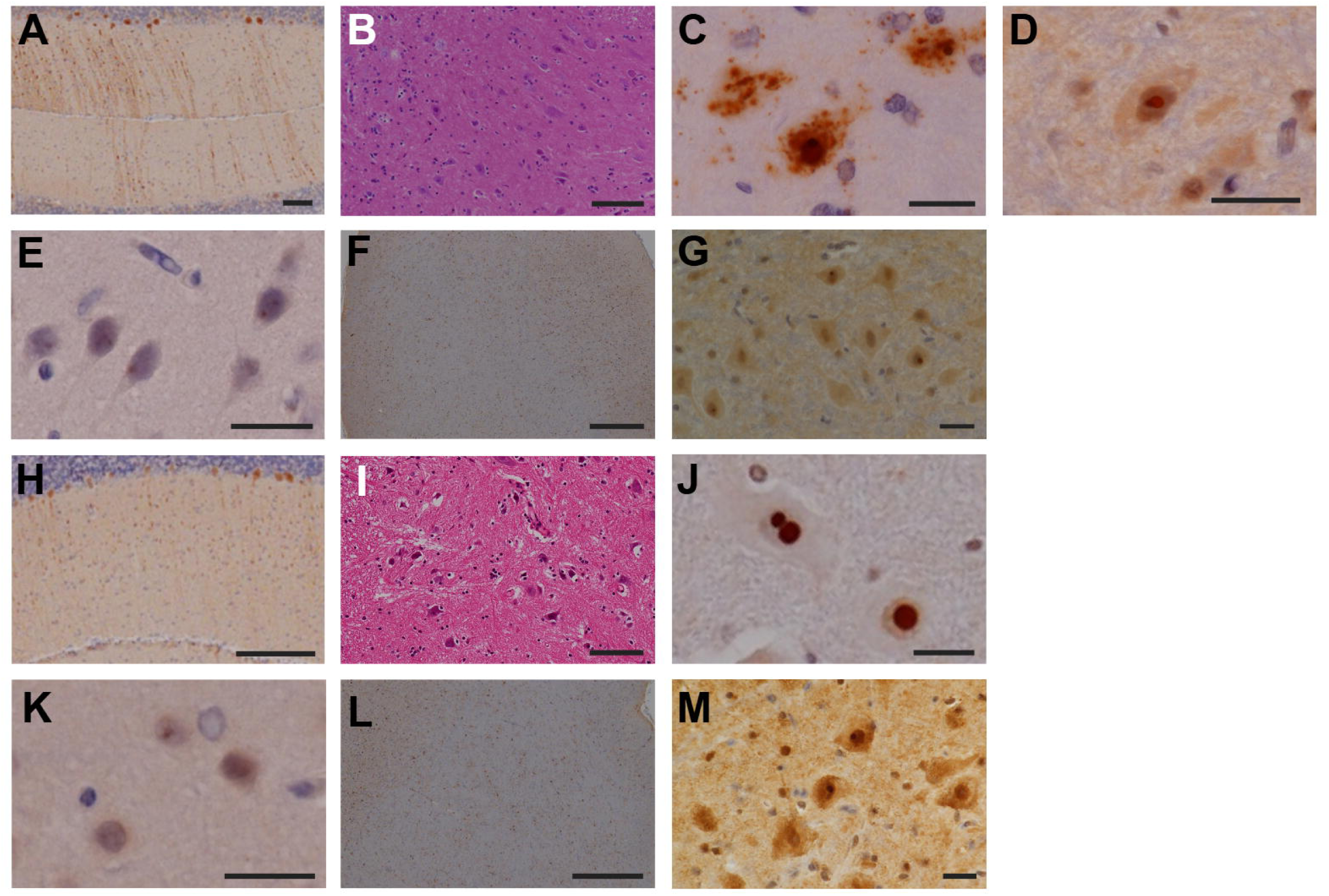
Degeneration and inclusion body formation in the brains, spinal cords, peripheral nerves and skeletal muscles of PQD13 and 14. A and. **H.** Anti-calbindin staining of the cerebellum of PQD13 (**A**) and PQD14 (**H**). **B–D, I and J.** Hematoxylin and eosin (**B** and **I**), anti-1C2 (**C**), and anti-ubiquitin (**D** and **J**) staining of the lumbar spinal cords of PQD13 (**B–D**) and PQD14 (**I** and **J**). **E and K.** Anti-ubiquitin staining of the cerebrum of PQD13 (**E**) and PQD14 (**K**). **F, G, L and M.** Anti-Iba1 (**F** and **L**) and anti-ubiquitin (**G** and **M**) staining of the brainstem of PQD13 (**F** and **L**) and PQD14 (**G** and **M**). Scale bars: 100 μm in A, B, and I; 20 μm in C–E, G, J, K, and M; 500 μm in F and L; 200 μm in H.

Altogether, these results indicate that the pathological findings from the founder marmoset (PQD1) were recapitulated in the symptomatic second-generation marmosets (Table 2). In addition, PQD13 and 14, which exhibited similar insertion sites and transgene copy number (Fig. 1) as well as comparable physical and behavioral phenotypes with early onset and rapid progression (Fig. 2 and 3), showed similar pathological phenotypes. These findings strongly suggest that the transgene causes the phenotype of these animals and indicate a probable mechanistic link between the transgene and the observed *in vivo* and post-mortem phenotypes.

### Brain region-specific characterization of transgene expression levels

Finally, we examined whether the differences in CNS transgene expression levels among the second-generation marmosets resulted in the aforementioned variability in behavioral and pathological changes. Although transgene expression in skin-derived fibroblasts was comparable among PQD11, 13, and 14 (Fig. 1D), we observed significant inter-animal variability in transgene expression in the CNS (Fig. 6). The transgene expression levels of both PQD13 and 14 were significantly higher than those of PQD11 in the thoracic and lumbar spinal cords, striatum, and temporal cerebral cortex, and appeared to have a higher tendency in the cerebellum (Fig. 6). These findings suggests that differences in transgene expression levels in these regions may contribute to differences in the severity of physical and behavioral impairments in these marmosets. This concept is compatible with the observed changes in patients with SCA3, in whom progressive motor impairments result from neuronal dysfunction in CNS regions including the cerebellum, spinal cord, and striatum (McLoughlin et al., 2020; Rüb et al., 2013). Together, these findings suggest the validity of our marmosets as a model of SCA3. In addition, the transgene expression levels were significantly lower in the frontal and occipital cerebral cortex and brainstem of PQD13 and 14 than of PQD11 (Fig. 6). This suggests that the neurotoxicity caused by transgene expression in these regions may have a minimal contribution to the differences in severity of the physical and behavioral impairments in these marmosets, which is also compatible with findings from patients with SCA3 suggesting that the cerebral cortex is affected to a lesser extent than the cerebellum (Yamada et al., 2001). Furthermore, the inter-animal variability of pathological changes in our marmosets (Table 2) was not associated with transgene expression levels or the severity of physical and behavioral impairments, except in the brainstem, where the difference in transgene expression level was associated with the presence of neurodegeneration in PQD11 but not in PQD13 or 14 (Table 2). We also observed different transgene expression levels between the two early-onset marmosets in several CNS regions; PQD13 showed significantly higher expression levels than PQD14 in the hippocampus, striatum, and cervical spinal cord, and significantly lower expression than PQD14 in the lumbar spinal cord, although the transgene integration site and copy number were equivalent in these animals (Fig. 1E, F). Overall, the severity of behavioral phenotypes in the second-generation marmosets (Figs. 2 and 3) correlated more strongly with transgene expression levels than with the neurodegenerative changes or inclusion pathology (Fig. 4, 5, and Table 1) in the CNS regions in which neuronal dysfunction is present in patients with SCA3 (Fig. 6). These findings suggest that neuronal dysfunction caused by transgene expression, rather than the resulting neuronal cell death, in these CNS regions may be the primary determinant of behavioral phenotypes in these animals.

**Figure 6.**
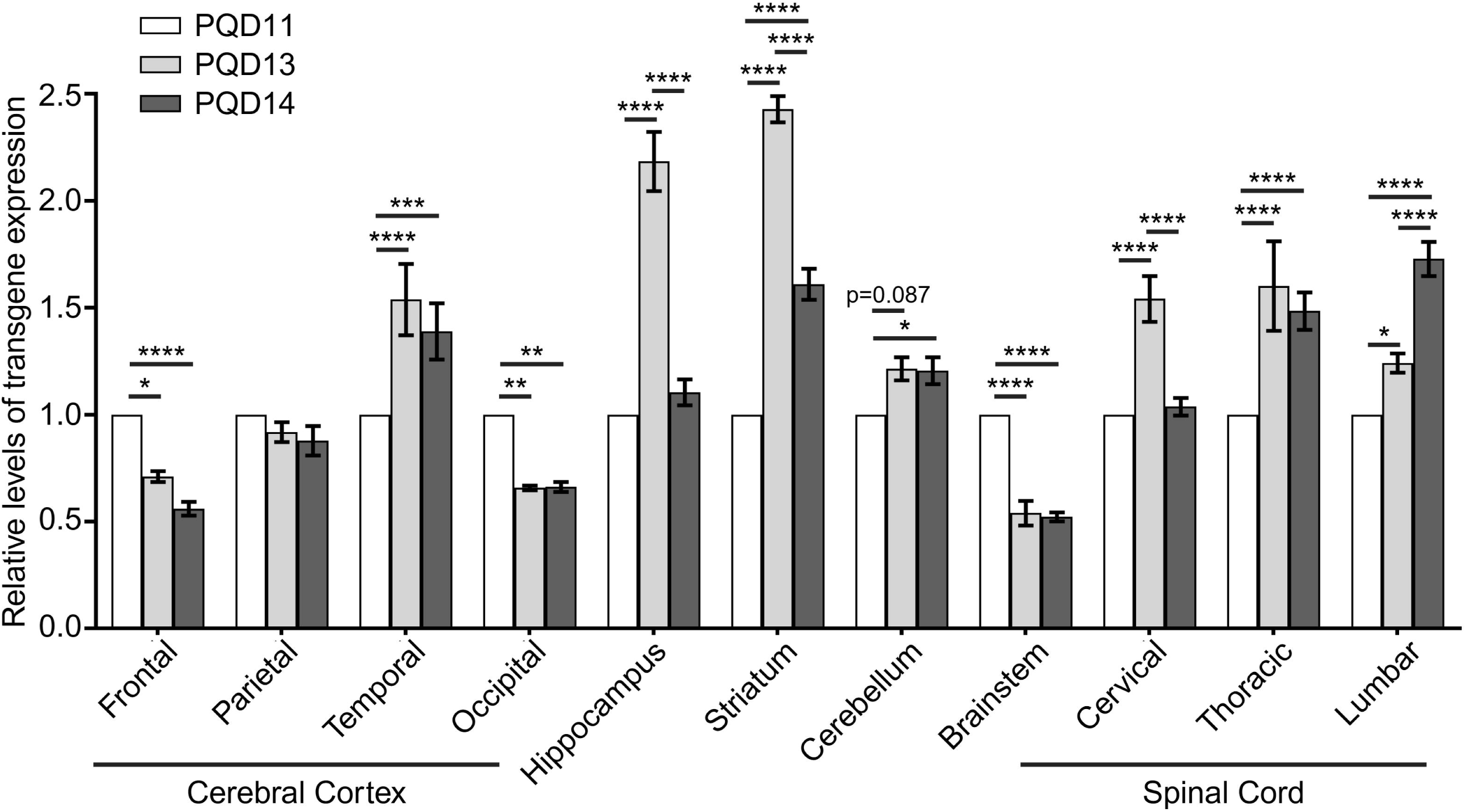
Comparison of region-specific transgene expression levels among symptomatic second-generation marmosets. Bar graph showing the comparison of relative transgene expression levels among regions and symptomatic marmosets. Transgene expression was normalized using GAPDH, and the normalized transgene expression level of PQD11 in each region was set as 1. Data are presented as the mean ± s.e.m. of four independent experiments. Statistical analyses were performed to assess inter-animal differences between the relative transgene expression levels using two-way ANOVA followed by Tukey’s multiple comparison test (* p < 0.05, ** p < 0.01, *** p < 0.001, **** p < 0.0001).

## Discussion

In the present study, we compared the genetic, behavioral, and pathological phenotypes between the founder and second-generation offspring of a transgenic marmoset model of SCA3. The transgenes were stably transmitted from the founder to all second-generation marmosets. Behavioral phenotypes, such as motor impairments and decreased weight gain and grip strength, and pathological phenotypes, such as a loss of Purkinje cells and an accumulation of polyQ proteins, were transmitted to three of the five second-generation marmosets. Our results indicate that we successfully generated a second-generation transgenic marmoset model of SCA3 with stable and reproducible phenotypes from the founder generation. Considering that the genetic mosaicism of the founder marmoset is eliminated in the second-generation marmosets, these second-generation marmosets will provide a valuable resource for future research once the colony is expanded.

### Advantages of SCA3 marmosets over previous NHP models of human diseases

Compared with rodent models, gene modification in NHPs has been envisioned as a way to create human disease models that better genocopy and phenocopy human diseases (Schmidt et al., 2022). The first human disease model that used genetically engineered NHPs was generated in rhesus monkeys to recapitulate Huntington’s disease (HD), another polyQ diseases, by overexpressing mutant *HTT* with expanded CAG repeats (Yang et al., 2008). In the founder animals, behavioral phenotypes resembling patients with HD, such as dystonia and chorea, and pathological hallmarks of HD in the brain, such as inclusions and aggregates immunoreactive for expanded polyQ protein, were observed (Yang et al., 2008). However, germline transmission of *HTT* was not achieved because neither of these founder animals survived until sexual maturity. The same group later generated a different founder animal harboring mutant *HTT* in the sperm and succeeded in obtaining two offsprings harboring mutant *HTT* (Moran et al., 2015). However, the CAG repeat length of mutant *HTT* was shorter in one of the offsprings (Moran et al., 2015) because of variations in CAG repeat length in the sperm DNA of the founder animal (Clever et al., 2019); this is also a well-known phenomenon also in humans because of the instability of triplet repeat sequences during DNA replication (Lenzmeier and Freudenreich, 2003; Wheeler et al., 2007). Given that the severity of the behavioral phenotypes in these animals correlated with the CAG repeat length, similar to human HD patients (Yang et al., 2008), the CAG repeat instability in these animals would likely lead to interindividual variation of behavioral phenotypes in the offsprings. This would be a limitation if the colony were expanded to study the underlying mechanisms of HD or develop novel treatments in the future.

Another human disease model using transgenic cynomolgus monkeys was generated to recapitulate Rett syndrome, a severe developmental disorder with autistic phenotypes, by overexpressing human *MECP2* (Liu et al., 2016). The founder animals showed brain-specific *MECP2* expression under the synapsin promoter and behavioral phenotypes that resembled patients with Rett syndrome, such as repetitive movements, anxiety, impaired social interaction, and mild cognitive dysfunction (Liu et al., 2016). Importantly, transgene expression and impaired social interaction were confirmed in the offspring of the founder animal (Liu et al., 2016). However, there was no correlation between the transgene copy number and the extent of behavioral abnormalities in these animals (Liu et al., 2016).

Our SCA3 model was the first human disease model generated in common marmosets (Tomioka et al., 2017). We previously reported that the founder animals showed behavioral and pathological abnormalities that resembled patients with SCA3, and that four offspring expressing the transgene were obtained from the founder marmoset (Tomioka et al., 2017). In the present study, we obtained one additional second-generation SCA3 marmoset expressing the transgene (Fig. 1B, D) and demonstrated that three of the five offspring exhibited behavioral and pathological phenotypes that resembled both the founder marmoset and patients with SCA (Figs. 2–5). Importantly, all five offspring exhibited similar CAG repeat length to the founder marmoset (Fig. 1C). This finding indicates that we successfully eliminated the CAG repeat instability by introducing one CAA triplet, which also encodes glutamine, in every 30 CAG repeats of the genetic construct that was used to establish the founder marmoset (Tomioka et al., 2017). In addition, we revealed that the severity of behavioral phenotypes in the symptomatic second-generation marmosets correlated with transgene expression levels in brain regions similar to those affected in patients with SCA3 (Figs. 2, 3, and 6).

These advantages make our SCA3 marmoset model suitable for future colony expansion, to allow the further investigation of underlying disease mechanisms or the development of novel therapeutics for SCA3 by other researchers, because phenotype consistency among animals is critical for such studies.

### Genetic determinants of the behavioral phenotypes of SCA3 marmosets

We previously reported that, among the symptomatic founder marmosets, phenotypic severity correlated with transgene expression levels in skin-derived fibroblasts (Tomioka et al., 2017). In addition, these transgene expression levels correlated with the number of transgene integration sites in the fibroblasts (Tomioka et al., 2017). In this study, we first demonstrated that transgene expression levels in skin-derived fibroblasts of the second-generation marmosets were similar among all symptomatic marmosets (Fig. 1D; PQD11, 13, and 14) despite variations in their behavioral phenotypes, which was in clear contrast with the founder generation. Subsequently, when we examined transgene expression levels in the spinal cord, striatum, and cerebellum, the CNS regions affected in patients with SCA3 exhibiting a progressive motor impairment (McLoughlin et al., 2020; Rüb et al., 2013), we demonstrated that expression levels were significantly higher in the marmosets with the early onset and rapid progression (PQD13 and 14) than in the marmosets with late onset and slow progression (PQD11). Importantly, these differences in cerebrospinal transgene expression did not correlate with the severity of cerebrospinal degeneration or inclusion formation changes (Table 2) at the endpoint, when all symptomatic marmosets reached a similar severity of behavioral phenotypes (Figs. 2A, B). Our observations therefore suggest that, even at the endpoint when motor impairment has nearly plateaued, transgene expression levels in the CNS still reflect the earlier disease course, such as the onset and progression of motor impairment.

The number of transgene integration sites (Fig. 1E) and the transgene copy number (Fig. 1F) were higher in the late-onset and slow-progression marmoset (PQD11) than in the early-onset and rapid-progression marmosets (PQD13 and 14), and were discordant with transgene expression levels in the fibroblasts or CNS of these marmosets (Figs. 1D and 6). These differences suggest that some of the transgenes were silenced by chromosomal position effects after being randomly integrated into the marmoset genome by lentiviral vector (Voigt et al., 2008). The positional effect may also have altered transgene expression in some of the CNS regions and resulted in the differences in transgene expression between skin fibroblasts and the CNS in symptomatic marmosets (Voigt et al., 2008). In addition, because congenic breeding is not performed in marmosets, differences in genetic backgrounds among the second-generation marmosets may have resulted in additional positional effects, thereby causing increased variability in the transgene expression levels, especially those observed in the CNS of the littermates, PQD13 and 14. Whole genome sequencing of all second-generation marmosets, longitudinal monitoring of the behavioral phenotypes in PQD12 and 15 for detecting later symptom onset, and pathological analyses of PQD12 and 15 may help to clarify the causes of phenotypic variability among second-generation marmosets as well as the genetic determinants of their behavioral phenotypes.

### Better validity of second-generation SCA3 marmosets as human disease models

Our second-generation marmosets offer better validity as an animal model of SCA3 than the founder marmosets. Three criteria are often used to validate animal models of human diseases, namely construct validity, face validity, and predictive validity (Tadenev and Burgess, 2019).

Construct validity is the degree of similarity between the mechanisms underlying the phenotypes of a certain animal model and those underlying the symptoms and pathology in humans; it is considered the most important criterion for validating animal models (van der Staay et al., 2009).

The construct validity of polyQ animal models is generally high because polyQ diseases are monogenic diseases with high penetrance caused by disease-causing expansions of CAG repeats encoding glutamine (Figiel et al., 2012; Lieberman et al., 2019). Our second-generation marmosets offer better construct validity than our founder marmosets because the genetic mosaicism of the founder was eliminated and there were fewer transgene integration sites in the second-generation marmosets (Fig. 1E).

Face validity is the degree of similarity in behavioral and pathological impairments between animal models and patients affected by the disorder (van der Staay et al., 2009). In contrast with the acute onset and rapid progression of motor impairments observed in our founder marmoset (Tomioka et al., 2017), one of our a second-generation marmosets, PQD11, exhibited a milder onset and progression of motor impairments that better recapitulated the disease course of patients with SCA3; it also exhibited a pathological phenotype that was similar to that of patients with SCA3. In human patients, SCA3 pathology is not limited to the CNS; peripheral neuropathy affects up to 60% of patients with SCA3 (C. França Jr. et al., 2009; Durr et al., 1996; McLoughlin et al., 2020). In addition, chronic myopathic changes accompanied by nuclear polyQ inclusions in the muscle fibers have been observed in a patient with SCA3 (Sugiyama et al., 2019). Notably, these behavioral and pathological phenotypes were reproduced in second-generation marmosets that harbored different genetic backgrounds from the founder marmoset, thereby resembling human conditions more closely than congenic rodent models (Vallender and Miller, 2013). In addition, we were able to test face validity in a reliable and reproducible manner by using MBAS, which we developed for longitudinal motor assessment in marmosets with less confounding factors than task-based motor assessments (Grieco et al., 2021; Verhave et al., 2009).

Predictive validity is defined by the ability of an animal model to predict the molecular and symptomatic drug responses in patients (Figiel et al., 2012). Results in the present study warrant colony expansion to test the predictive validity in future research. Although it is generally difficult to test predictive validity in diseases with poor therapeutic standards, including polyQ diseases where disease-modifying therapies remains a clinically unmet need (Figiel et al., 2012; van der Staay et al., 2009), the recent identification of candidate compound and antisense oligonucleotides for disease modification in these conditions may help to test the predictive validity of our SCA3 model marmosets (Ishihara et al., 2024; Minakawa et al., 2020; Minakawa and Nagai, 2021; Moore et al., 2019, 2017). In addition, our successful generation of a transgenic marmoset disease model with abnormal protein accumulation and neurodegeneration paves the way for the establishment of gene-modified marmoset models of other neurodegenerative diseases in general, not limited to SCA3.

This will then enhance the development of translatable biomarkers and disease-modifying therapies for these currently intractable diseases.

## Materials and Methods

### Experimental animals

All animal experiments were approved by the ethics committee for primate research of the National Center of Neurology and Psychiatry in Japan. All experiments were conducted in accordance with institutional guidelines and the National Research Council’s Guide for the Care and Use of Laboratory Animals.

Five second-generation transgenic common marmosets (*C. jacchus*), PQD 11–15 (three male and two female), were established from PQD1, one of the first-generation transgenic marmosets (Tomioka et al., 2017), and were used in the present study. PQD1, 11, 12, and 15 are male, and PQD 13 and 14 are female. Of the second-generation marmosets, PQD 11–14 were subjected to genotype analyses (Tomioka et al., 2017) and PQD11, 12, and 15 were subjected to in-cage motility assessments (Koizumi et al., 2021) in our previous studies. In addition, 11 (five male and six female) or five (two male and three female) wild-type marmosets were used as controls for the body weight measurements and bar grip test (described in the *Bar grip test* subsection), respectively. All animals were housed under standard conditions under a constant 12-h dark/12-h light cycle. Food was provided twice a day in the morning and the evening. Water was provided *ad libitum*. All animal experiments and housing conditions were approved by the committee of ethical issues in animal experiments of the National Center of Neurology and Psychiatry, Japan, and were performed in accordance with the institutional guidelines for the care and use of primates. The institutional guidelines complied with the guidelines issued by the Japanese Ministry of Education, Culture, Sports, Science and Technology and the Guide for the Care and Use of Laboratory Animals (National Research Council (US) Institute for Laboratory Animal Research, 1996).

### Production of second-generation transgenic marmoset

Second-generation marmosets were produced from PQD1 as previously described (Tomioka et al., 2017). Briefly, oocytes obtained from wild-type marmosets were fertilized with sperm from PQD1 via intracytoplasmic sperm injection. The embryos were then cultured and transferred to surrogate mothers. After embryo transfer, the surrogate mothers were monitored monthly for pregnancy by ultrasound imaging until delivery.

### Establishment of ear fibroblast cell lines from the transgenic marmosets

Ear fibroblast cell lines from transgenic marmosets were established and cultured as previously described (Tomioka et al., 2017).

### Genomic PCR analysis and reverse-transcription PCR analysis

Genomic DNA or RNA was extracted from ear fibroblast cell lines and first-strand cDNA was synthesized from the extracted RNA as previously described (Tomioka et al., 2017). Genomic DNA and cDNA were then subjected to PCR to detect mutant *ATXN3* or *ACTB* as previously described (Tomioka et al., 2017). The plasmids used to create the first-generation transgenic marmoset were used as positive controls.

### CAG repeat analysis

To detect CAG repeats within the integrated transgene of each marmoset, cDNA generated from ear fibroblast cell lines was subjected to PCR as previously described (Tomioka et al., 2017). To detect *ACTB*, the *ACTB* forward 1 and reverse 2 primers (Tomioka et al., 2017) were used. The original plasmids used to create the first-generation transgenic marmoset were used as positive controls.

### Quantitative reverse transcription PCR (qPCR) analysis

The relative expression levels of the integrated transgenes were determined using qPCR analysis with cDNA generated from ear fibroblast cell lines or post-mortem brains as templates; this was performed as previously described with slight modification (Tomioka et al., 2017). The primers used to detect the expression level of the transgene were CAG forward 2 (5′-ACTTGGGAGTGATCTAGGTGATG-3′) and HA reverse (5′-AGTCGGGGACGTCGTAGGGGTATT-3′). All qPCR results were normalized using *GAPDH* as previously described.

### Integration site analysis of the transgene

The integration sites of the lentiviral vector harboring the transgene were determined using the LentiX integration site analysis kit (Takara Bio, Kusatsu, Japan) according to the manufacturer’s instructions. The integration sites were confirmed by subjecting genomic DNA extracted from ear fibroblast cell lines to PCR for 35 cycles of 98°C for 10 s and 68°C for 60 s using Emerald Amp PCR Master Mix (Takara Bio, Kusatsu, Japan) with the following primers: Chr1 forward (5′-TTTCCCTGCTGGCTGCTCTACTGC-3′), Chr3 forward (5′-AGTGCATATCCCGACACCTTGCTGG-3′), Chr4 forward (5′-AGATGGTGCCCACCCAGATTGAGG-3′), Chr10a forward (5′-GAGATGGAGTCTCACGGTGTTGCCTAG-3′), Chr10b forward (5′-GGTAGTGTGAAAGAGGCCCTGAAGC-3′), Chr16 forward (5′-AT GATCTGCCCGTCTTGGCCTCC-3′), and LSP2 reverse (5′-GCTCCTCTGGTTTCCCTTTCGCTTTCAA-3′).

### Digital PCR analysis

Copy numbers of the integrated transgene in the second-generation transgenic marmosets were determined using digital PCR analysis. Genomic DNA extracted from ear fibroblast cell line was digested using restriction enzyme (BamHI) and subjected to digital PCR for 39 cycles of 60°C for 120 s and 98°C for 30 s using QuantStudio 3D dPCR Master Mix (Applied Biosystems, Waltham, USA.) according to manufacturer’s instructions using the following primers: ATXN3 (5′-FAM-CAGGCTCCGCGGCCGCCGTA-3′) and NR0B1 (5′-VIC-CGGGTCGCAGCGCACCACAG-3′) probes and CMV-Fwd (5′-GTGGATAGCGGTTTGACTCACG-3′), ATXN3-Rev (5′-TGTTGAGCACAAAGTGAGCCTTC-3′), NR0B1-Fwd (5′-CAGGCCATGGC TCTTCTGTA-3′) and NR0B1-Rev (5′-GGCTCTGAGGTTTCCACAGT-3′). All digital PCR results were normalized using NR0B1.

### Semiquantitative rating scale for behavioral abnormality assessment

We developed a semiquantitative rating scale, the MBAS, to assess behavioral abnormalities of the marmosets using a 10-minute-long video recorded in their home cage and evaluated the onset and progression of the motor impairments in the second-generation transgenic marmosets. The MBAS included five categories: posture, activity, movement, muscle strength, and balance (Table 1).

Marmosets were rated from 0 to 3 points in each of these categories (0 point = normal, 1 point = mildly abnormal, 2 points = moderately abnormal, and 3 points = severely abnormal; maximum total score = 15 points) based on the video, which was recorded using a standard video camera placed in front of the home cage between 2:00 p.m. and 4:00 p.m. After all videos were recorded, the assessments were performed by the rater (N.N.), who was blinded to the timing of each recording but not to the individual animals, because each animal was able to be visually identified by their facial features in the videos).

### Quantitative in-cage motility measurement

In-cage marmoset motility was measured using an in-cage motility monitoring method for new world monkeys as previously described (Koizumi et al., 2021). Briefly, an infrared motion sensor was placed above the cage to detect the movement of each marmoset within their cage. The pulse counts detected using the infrared sensor were converted to a binary queue to generate a basic unit for accurately counting the number of movements of each marmoset. The marmosets received continuous 24-hour monitoring throughout the measurement period, and total activity count per 24 hours were calculated.

### Bar grip test

The neuromuscular functions of the upper extremities of each marmoset were evaluated using the bar grip test as previously described, with slight modifications (Tomioka et al., 2017). Briefly, the grip force of each marmoset was recorded by letting the marmoset grasp the bar (MK-380-M, Muromachi Kikai, Kyoto, Japan) with their maximal grip force. The maximal grip force was consecutively measured five times and the average was recorded.

### Histopathological analyses

Symptomatic marmosets were subjected to pathological analyses at the age of 15 months for PQD11 and 12 months for PQD13 and 14. Histopathological analyses of the postmortem brains, spinal cords, peripheral nerves from the upper and lower extremities, and quadriceps femoris muscle of PQD13 and 14 were performed as previously described (Tomioka et al., 2017). Briefly, the tissues were fixed with 4% paraformaldehyde, embedded in paraffin, and sectioned at 5 μm thickness. Hematoxylin and eosin staining was performed using standard procedures. Immunohistochemical staining was performed using the following primary antibodies: monoclonal anti-1C2 (1:5000, Merck Millipore), polyclonal anti-ubiquitin (1:1000, Dako), monoclonal anti-GFAP (1:500, Agilent), polyclonal anti Iba-1 (1:1000, Wako), and polyclonal anti-calbindin (1:16,000, Swant). Peripheral nerves were stained with toluidine blue. Lumber anterior horn were prepared for electron microscopy. These tissues were post-fixed in 2.5% glutaraldehyde in phosphate buffer (pH 7.4), osmicated, and then processed for embedding in plastic resin. Semithin 1 μm sections were stained with toluidine blue for light microscopic observation. Appropriate areas were selected and trimmed. Ultrathin sections were stained with uranyl acetate and lead citrate and examined with an electron microscope.

### Statistical analyses

All statistical analyses were performed using GraphPad Prism 9. *P* < 0.05 in two-tailed analyses was considered to indicate a statistically significant difference.

## Acknowledgements

The authors are grateful to Drs. Yuji Takahashi and Yuji Saito (Department of Neurology, National Center Hospital, National Center of Neurology and Psychiatry) and Dr. Keiji Wada (National Center of Neurology and Psychiatry) for their valuable discussions and suggestions on this study. We also thank Bronwen Gardner, PhD, from Edanz for providing professional English editing of this manuscript.

## Competing interests

All authors declare that the research was conducted in the absence of any financial, personal or professional associations that could be perceived as a potential conflict of interest.

## Funding

This work was supported by Intramural Research Grants for Neurological and Psychiatric Disorders from the National Center of Neurology and Psychiatry [26-11, 29-5, and 2-8], the Japan Agency for Medical Research and Development [JP16ak0101027, JP19ek0109216, JP23ek0109522, and 26-005], and Grant-in-Aid for Young Scientists (B), JSPS KAKENHI Grant Number 26870884.

## Data availability

The data that support the findings of this study are available from the corresponding authors, upon reasonable request.

## Author contributions statement

Conceptualization: K.S., I.T.

Methodology: I.T., K.O., N.N., M.K., R.S., Y.K., T.N., A.K.

Investigation: E.N.M., K.O., N.N., T.N., M.K., A.K., Yo.S., M.K., S.N., T.H., Y.H., H.N., T.T., Y.K.,

R.S., Y.N., Yu.S., I.T., K.S.

Validation: E.N.M., T.N.

Formal analysis: E.N.M., N.N., T.N., K.O.

Writing – original draft preparation: E.N.M.

Writing – review and editing: K.S., T.N., Yu.S., T.T., Y.N.

Visualization: E.N.M., K.O., N.N., T.N.

Supervision: K.S., I.T., Y.N.

Project administration: K.S.

Funding acquisition: K.S., I.T.

## Notes

### Competing Interest Statement

The authors have declared no competing interest.

